# Inter-individual differences in human brain structure and morphometry link to population variation in demographics and behavior

**DOI:** 10.1101/413104

**Authors:** A. Llera, T. Wolfers, P. Mulders, C. F. Beckmann

**Affiliations:** Donders Institute for Brain, Cognition and Behavior, Radboud University, Nijmegen, The Netherlands.; Donders Centre for Cognitive Neuroimaging, Nijmegen, The Netherlands.; Department of Cognitive Neuroscience, Radboud University Medical Centre, Nijmegen, The Netherlands.; Oxford Centre for Functional Magnetic Resonance Imaging of the Brain (FMRIB), Oxford University

## Abstract

We perform a comprehensive integrative analysis of multiple structural MR-based brain features and find strong evidence relating inter-individual structural variations to demographic and behavioral variates across a large cohort of healthy human volunteers. In particular, our findings shed some light on functional & structural integration, as we find a mode of structural variation that relates to and extends the ‘positive-negative’ behavioral spectrum which was recently reported as being associated with variations in functional connectivity.

**Significance statement:** This work provides for the first-time strong evidence relating human brain structure variations to a *wide range* of demographic and behavioral measures. We show that several measures previously associated to variation in functional MRI connectivity are in fact already associated at the structural level, pointing towards structure-function integration.

## Introduction

Understanding individual human behavior has attracted the attention of scientists and philosophers since antiquity. The first quantitative approach intended to deepen such understanding dates to the first half of the 19-th century when ‘phrenology’^1,2^ related human behavior or cognitive abilities to skull measures. Technical, intellectual and clinical advances in the last two centuries allow us to now accurately quantify brain structure and function^3–7^, and to summarize certain ‘aspects’ of human behavior by means of standardized tests. Such advances facilitate automatized exploratory statistical learning analyses to uncover previously hidden relationships between brain features and human behavior, demographics or pathologies^8^. These developments are expected to be pushed even further with the emergence of the big data MR imaging epidemiology phenomenon^9,10^, and some exponents of such expectations have already reported associations with blood-oxygen-level dependent (BOLD) brain function^11,12^; for example, functional connectivity patterns can be used to identify individuals^11^, predict fluid intelligence^11^, or describe a mode of functional connectivity variation that relates to lifestyle, happiness and well-being^12^. Although the brain’s function-structure relationships are not yet fully understood, linking structure to behavior is essential for either type of imaging modality to be fully interpretable as an imaging phenotype. Further, given the long-term character of some demographic variables (e.g. overall happiness), we hypothesize that different brain structural features such as regional variation in the density of gray matter or subject-dependent degree of cortical expansion should also, to some extent, reflect these relationships with behavior. To test these hypotheses, in this work we make use of the large quantity of high quality behavioral and neuroimaging data collected by one of the big data initiatives, the Human Connectome Project ^9^ (HCP). The HCP sample includes detailed structural imaging, diffusion MRI, resting-state and several different fMRI tasks for each subject. Furthermore, the availability of more than 300 behavioral and demographic measures allows the post-hoc exploration of a wide range of associations^13^. We further hypothesize that behavioral variations can be explained by more general brain structure variations than isolated single feature variations (e.g. cortical thickness variations); we consequently extract multiple structural features from the different MR modalities and perform a simultaneous analysis by linked independent component analysis (Linked ICA^13,14^). Such analyses increase statistical power by evidence integration across different features^15,16^ and have been shown to be powerful in identifying correlated patterns of structural and diffusion spatial variation that can then be studied in relation to individual behavioral and demographic measures^17,18,16^. Although similar analyses have been previously performed^17^, in this work we benefit from the unique characteristics of the data sample; we consider brain and behavioral data from close to 500 “healthy young adults” which reduces common pathology- and age-related variance and so increases the power to detect associations due to normal cross-sectional variability.

Our results support the hypothesis that structural brain features are strongly associated with demographic and behavioral variates. Interestingly, the most relevant mode of structural variation identified through the multi-modal data fusion approach relates to recent findings obtained using functional MRI data from the same HCP cohort. In particular, our findings closely resemble the ‘positive-negative’ set of behavioral measures identified in Smith et al.^12^ on the basis of functional (co-)variations, pointing towards structure-function integration. Indeed, they support the notion that it is the spatial configuration in functional parcels rather than cross-parcel connectivity^19^ that is driving the brain-behavior relationships.

## Methods

In this work, we use data from the Human Connectome Project (HCP) N=500 release which contains data from healthy young adults including twins and their non-twin siblings. In addition to performing more than 300 behavioral/demographic tests, each subject participated in structural, diffusion and several functional MRI recordings^20,21^. A description of all MRI and behavioral/demographic measures included in our analysis can be found in van Essen et al.^20^ and we provide a summary of the latter in the supplemental information (SI), Table S1. Due to structure-function integration we hypothesize that different biological features such as regional variation in the density of gray matter, white matter connectivity or subject dependent degree of cortical expansion should reflect similar associations with behavior as the ones reported at ^11,12^. To investigate such hypothesis, the structural MRI T1-weighted images were used to extract gray matter densities and cortical measures, using a Voxel Based Morphometry (VBM)^22,23^ (http://www.fil.ion.ucl.ac.uk/spm12) pipeline to extract cortical gray matter probability maps as well as maps of cortical thickness (CT) and pial area (PA)^24,25^ estimates by means of transforming all anatomical T1-weighted cortical surfaces through FreeSurfer v5.3 (http://surfer.nmr.mgh.harvard.edu). Further, the diffusion-weighted MRI data were used to extract several features, i.e. fractional anisotropy (FA), anisotropy mode (MO) and mean diffusivity (MD) ^26,27^ (https://fsl.fmrib.ox.ac.uk/fsl/v5.0.9). In addition to these structural readouts and in order to also include purely local morphometric differences across subjects, we also consider the images containing the Jacobian determinants (JD) of the warp fields defining the transformations of each subject’s structural image onto a reference brain. These feature extraction operations are schematically summarized in Figure 1 operation A, and full details on the data processing performed to achieve each feature are provided in the SI under the section ‘*Individual features pre-processing’*. From the initial N=500 participants, several subjects were excluded on the basis of abnormalities in any of the features. In total, N=448 subjects were entered into further analyses. We then use the Linked-ICA model^13^ to simultaneous factorize the considered N=448 subjects’ VBM, FA, MO, MD, CT, PA and JD features into independent sources (or components) of spatial variation. In brief, Linked-ICA is an extension of Bayesian ICA^28^ to multiple input sets, where all individual ICA factorizations are linked through a shared common mixing matrix that reflect the subject-wise contribution to each component. This operation is represented in Figure 1 operation B where we can also appreciate that such factorization provides - per component - a set of spatial maps (one per feature modality), a vector of feature loadings that describe the degree to which the component is ‘driven’ by the different modalities, and a vector that describes how each individual subject contributes to a given component. Importantly, the subject-loadings define the cross-subject variation of the multi-modal effects and can subsequently be used to study relationships to other behavioral or demographic cross-subject variations by means of simple correlations. Given our sample size and following^13,14^ we report full results from a 100 dimensional factorization. Different model order decompositions were also performed to demonstrate the robustness to the choice of dimensionality; further, we also performed an analogous multi-modal analyses where we excluded the JD features as well as an independent component analyses^29^ of the JD features in isolation to evaluate the dependency of the results on purely morphometric differences-these are reported in the SI. To perform the multi-modal analyses we developed a new implementation of the Linked ICA algorithm^13^ which will be made publicly available as part of the next release of the FSL toolbox^27^.

**Figure 1:**
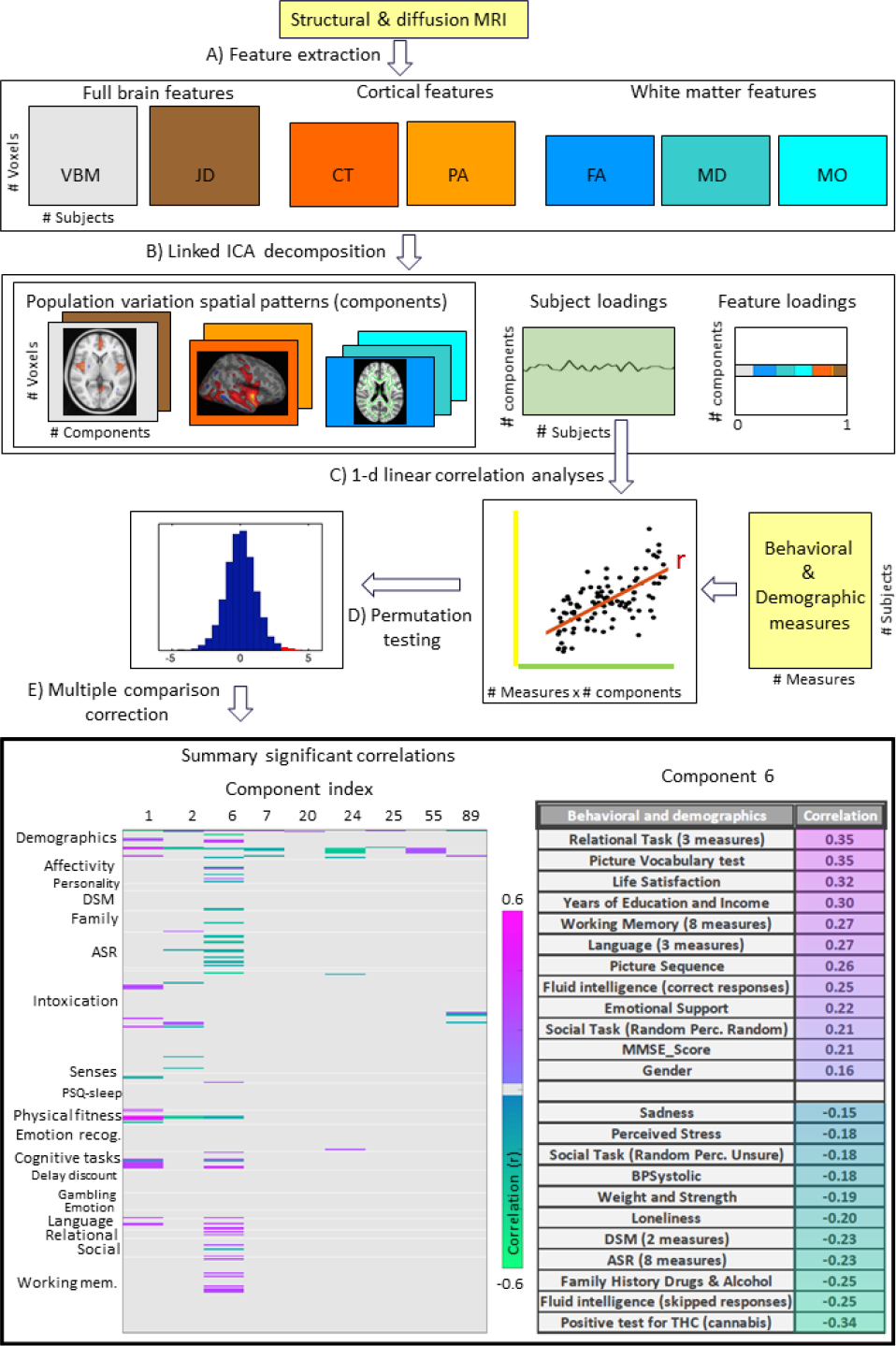
Data processing pipeline and main results. (A) Structural and diffusion-weighted MRI data are used to extract relevant features i.e. Voxel-Based Morphometry (VBM), Fractional Anisotropy (FA), Mean Diffusivity (MD), Anisotropy Mode (MO), Cortical Thickness (CT), Pial Area (PA) and Jacobian Determinants (JD). (B) These features are used as input to the Linked ICA algorithm. (C) Subject loadings of each independent component are fed together with the behavioral/demographic measures into a correlation analysis. The bottom left panel presents demographic and behavioral measures grouped by categories (y-axis), and a representative set of components reflecting significant correlation with at least one behavioral measure (x-axis). The color scale encodes the Pearson correlation coefficient and only significant correlations are color coded. In the bottom right panel we present a summary of component number 6 significant correlations to behavioral and demographic variates, resampling a mode of structural variation that links to and extends the ‘positive-negative’ behavioral spectrum previously attributed to functional connectivity variations^12^.

## Statistical analysis

To uncover relationships between the behavioral/demographic measures and the components obtained from the Linked-ICA decomposition we perform a correlation analysis between each independent component subjects’ contribution and each available behavioral measure. This operation is schematically summarized in Figure 1 operation C. To take into account the family structure present in the HCP sample while assessing significance we use the Permutation Analysis of Linear Models (PALM) ^30,31^ and use 10^6^ permutations per tested correlation (Figure 1 operation D). We define significance at p < 0.05 and address the multiple comparison by applying FDR correction^32^ as well as full Bonferroni correction (Figure 1 operation E).

## Results

The multimodal brain data analyses resulted in a total of 100 collections of component maps, each of which can be represented as 7 spatial maps covering the gray-matter space (VBM), diffusion skeleton space (DTI, MD, MO), cortical vertex space (CT and PA) and a voxel-wise map of the Jacobian deformation (JD). In addition, each collection of maps is associated with a single vector of contributions that describe the degree to which a given collection is ‘driven’ by the different modalities. Further, each collection is associated with a single vector that describes how each individual subject contributes to the component. Post-hoc linear correlation analyses of these subject contributions with behavioral measures identified, after FDR correction^1^, a total of 155 significant brain-behavior correlations, summarized by 30 components reflecting at least one significant relationship to behavior. We provide the full results in SI Table S2 and a brief summary in the bottom left panel of Figure 1 where we color code the significant Pearson correlation values for the components showing at least one Bonferroni corrected ^2^ significant correlation to a behavioral or demographic measure.

A single component, number 6, shows strong associations with 48 measures across several behavioral domains and across all structural modalities (SI Figure S2). In Figure 1 bottom right panel we provide a summary of the behavioral measures significantly correlating with component 6 as well as the corresponding Pearson correlation values. For interpretation, the behavioral measures are grouped and ordered according to a decreasing correlation value. Note that in the cases where several measures are grouped together we report their mean correlation value and full results can be again found in SI Table S2. We observe that component number 6 relates to various behavioral scores including working memory, language function and general wellbeing (life satisfaction, social support). In Figure 2 we present the associated spatial maps: VBM measures are most heavily weighted in bilateral orbitofrontal cortex, temporal pole, lingual gyrus and the putamen (first row). Morphometric differences (JD features) load into temporal lobes, caudate and brainstem (second row), and white matter tracts do most heavily weigh onto the internal capsule, anterior thalamic radiation and the anterior corona radiate (3^rd^, 4^th^, and 5^th^ rows). Cortical effects (6^th^ and 7^th^ rows) are largely associations with multi-modal association cortex that show effects whereas primary sensory cortices are not implicated. Note that the involvement of areas such as the putamen and lingual gyrus are relevant to explain the behavioral relationships found with working memory and word processing. Further, the involvement of structural connections between subcortical and prefrontal areas as well as the orbitofrontal cortex and temporal poles could explain the link to more complex functions such emotional support or life satisfaction.

**Figure 2:**
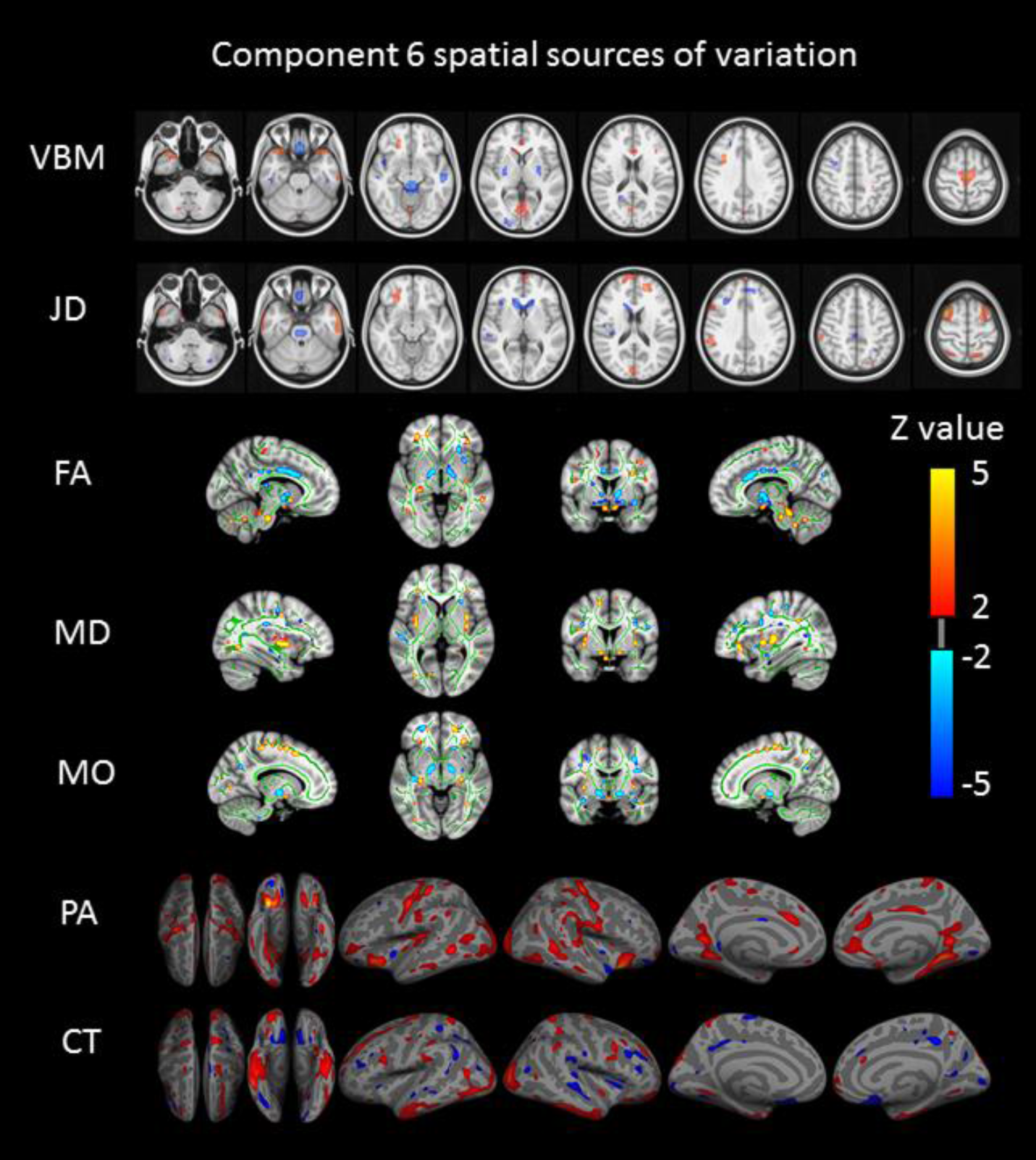
Component number 6 feature sources of variation. From top to bottom we visualize the VBM (Voxel Based Morphometry), JD (Jacobian Determinants), FA (Fractional Anisotropy), MD (Mean Diffusivity), MO (Mode of Anisotropy), PA (Pial Area), and CT (Cortical Thickness) spatial maps. For improved visualization, each modality has been thresholded at a z-value of 2. This mode of structural variation, component 6, that strongly reflects a ‘positive-negative’ behavioral spectrum, links to a wide range of brain regions across structural modalities and might reflect the structural multimodal foundation of a functional brain network linked to these variations that has been earlier identified.

Components 1 and 2 also show to be related to several measures: components 1 relates mainly to gender, physical strength and language and it is defined by significant changes in gray matter density (VBM measures) and cortical real expansion (PA measure). Similarly, component 2 is driven by VBM maps and correlates with variations in gender, age, height, weight and strength. Components 7, 24, 25, 29 and 55 are driven by at least 3 feature modalities and they map into gender, weight, body mass and height. Number 89 maps VBM and JD into hematocrit and component 20 maps VBM and JD into age. Details about the spatial extent of each of these components can be found in SI, section *‘spatial maps’*. Another set of components show associations, albeit either being associated with only a single modality and/or with a small set of behavioral variates (SI Table S2). Many of these components show simple relationships to overall size measures, such as weight, body mass (BMI) or height, and the associations are weaker than the above reported; we consequently decided to not further discuss their spatial extent in this work and we provide full nifty images in the SI.

To validate the robustness of the presented results to the model order choice we performed analyses at different dimensionalities and observe that especially lower indexed components are highly reproducible. In particular, component number 6 is recovered at dimensionalities 90 and 110 with a subject mode correlation value of around r∼0.9 (details can be found in the SI section *‘Robustness: model order’).* Regarding the influence of purely morphometric differences in the analyses, a comparative analysis excluding the JC revealed essentially unaltered brain-behavior associations. Further, an analyses of only the JD feature showed that no fully corrected significant association to the reported positive-negative structural mode is found when considering uniquely morphometric differences, even if considering several components together. However, uncorrected statistics suggests that information of the positive-negative mode could already be present at the morphometric level. These results are presented in SI section *‘Robustness: analyses without Jacobians’, and ‘Robustness: Analyzing morphometric differences’.*

## Discussion

We present a simultaneous analysis of brain structural measures that reveals how several types of behavior and demographics link to variations in such measures of brain structure. Several components detect simple associations between brain size (encoded in gray matter density and cortical area) being related to gender, strength, endurance or language function.

More interestingly, we encounter a single pattern of gray and white matter covariation that is strongly associated with several measures relating to cognitive function including working memory and language function, while also being strongly related to several measures of wellbeing including life satisfaction or emotional support. Accordingly, the spatial organization of the component that relates to these measures predominantly includes regions and connections that are relevant to working memory and word processing such as the putamen and lingual gyrus ^33,34^. Additionally, the inclusion of regions such as the orbitofrontal cortex and temporal poles, as well as structural connections from subcortical to prefrontal regions, could explain the link to more complex functions such as emotional support and life satisfaction. Furthermore, the mode of structural variation we report here relates to several recently reported results obtained using functional MRI. In particular, our results relate to the ones presented in Finn et al.^11^ since it identifies fluid intelligence measures and it also shares many behavioral measures also identified by the ‘positive-negative’ mode reported in Smith et al.^12^. Clearly, the functional analyses presented in Smith et al.^12^ and the one we just presented here, while using entirely different MRI measurements, are both able to get at the core of the same behavioral spectrum; our analyses reliably augments the spectrum of behavioral variables reported by the functional analyses by extending it with many working memory, language, relational task, ASR and DSM measures (Figure 1 bottom right and SI Table S2). It is to note here that while the statistics reported in Smith et al.^12^ were obtained from a Canonical Correlation Analyses (CCA) between partial correlation matrices and all behavioral measures at once, the statistics we present here involve simple linear correlations. While the former type of analysis can benefit from the multi-variate type of analysis through the application of CCA, ensuing results can be hard to interpret. The straight-forward individual linear correlation analysis against the behavioral/demographic measures separately instead affords simple interpretation, albeit possibly being over-conservative given the chosen significance level.

These findings directly look into the relationship between brain structure and function. In fact, the functional mode of variation is strongly associated with connectivity in brain areas approximately resembling the Default Mode Network and, given the spatial extent and the strong weight of the DWI data in the structural mode we report, it seems reasonable to assume that these white matter structure variations could contribute to the functional connectivity changes reported in Smith et al.^12^. Further we found no clear spatial overlap between the reported structural mode and the cortical functional extent of the ‘positive-negative’ mode, suggesting that functional-structural simultaneous analyses should increase the sensitivity of both, functional and structural analyses. Further, these results might question whether group functional connectivity measures using fMRI provide direct measures of brain connectivity or are biased due to individual structural differences that have been taken into account inadequately. An analogous multimodal analysis excluding the JD feature provided equivalent results to the presented here (SI Table S3) and uni-modal analysis of uniquely the JD features (using simple ICA-based decomposition^29^ of the single JD modality) did not provide significant correlation to the behavioral mode at the level of fully corrected statistics. These extra analyses confirm that the structural features relating to the behavioral mode are not uniquely driven by morphometric differences and consequently suggest that the behavioral associations found using functional MRI, although could be influenced, are not uniquely driven by the structural differences; these results seem to be aligned with recent findings by Bijsterbosch et al.^19^ where it is shown that individual spatial configurations extracted from functional MRI rather than the connectivity profiles between areas seem to stronger relate to the positive-negative mode.

In a future study, we will couple structural-functional modalities to investigate the link of these modalities more immediately. Here, we showed that a positive-negative mode of behavioral variation extends beyond the functional domain to also link to multimodal brain structure. This will have important implications in the future analyses of neuroimaging big data and will help improve our understanding over the functional-structural integration.

## Acknowledgements

We are grateful to Stephen M. Smith for the helpful discussions and to V. Kumar for help with the visualization of the DWI results. The research leading to these results has received funding through a Synergy Grant by the European Research Council under the European Union’s Seventh Framework Programme (FP/2007-2013), ERC Grant Agreement no. 319456. We further gratefully acknowledge support from the Netherlands Organization for Scientific Research (NWO) through VIDI grant to CFB (864.12.003) and we also gratefully acknowledge funding from the Wellcome Trust UK Strategic Award (098369/Z/12/Z).

## Footnotes

1 FDR corrected, q < 2.2×10^−4^.

2 Bonferroni corrected q < 1.4×10^−6^.

## Supplemental information (SI)

In Table S1 we provide a summary of the behavioral and demographic measures present in the Human Connectome Project (HCP) sample. For easier interpretation we grouped them here by categories and a full detailed description can be found in van Essen et al.^20^.

**Table S1:**
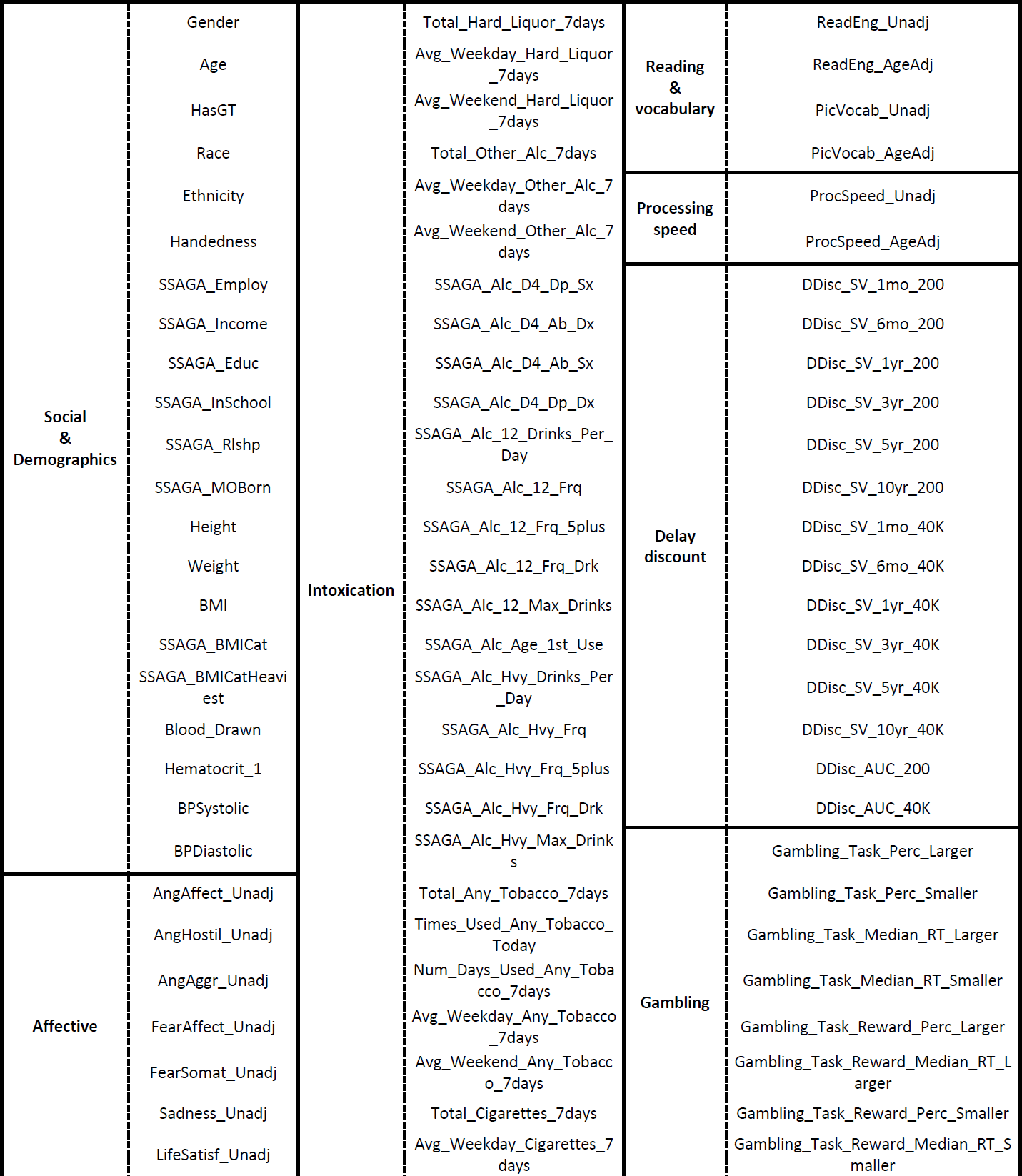
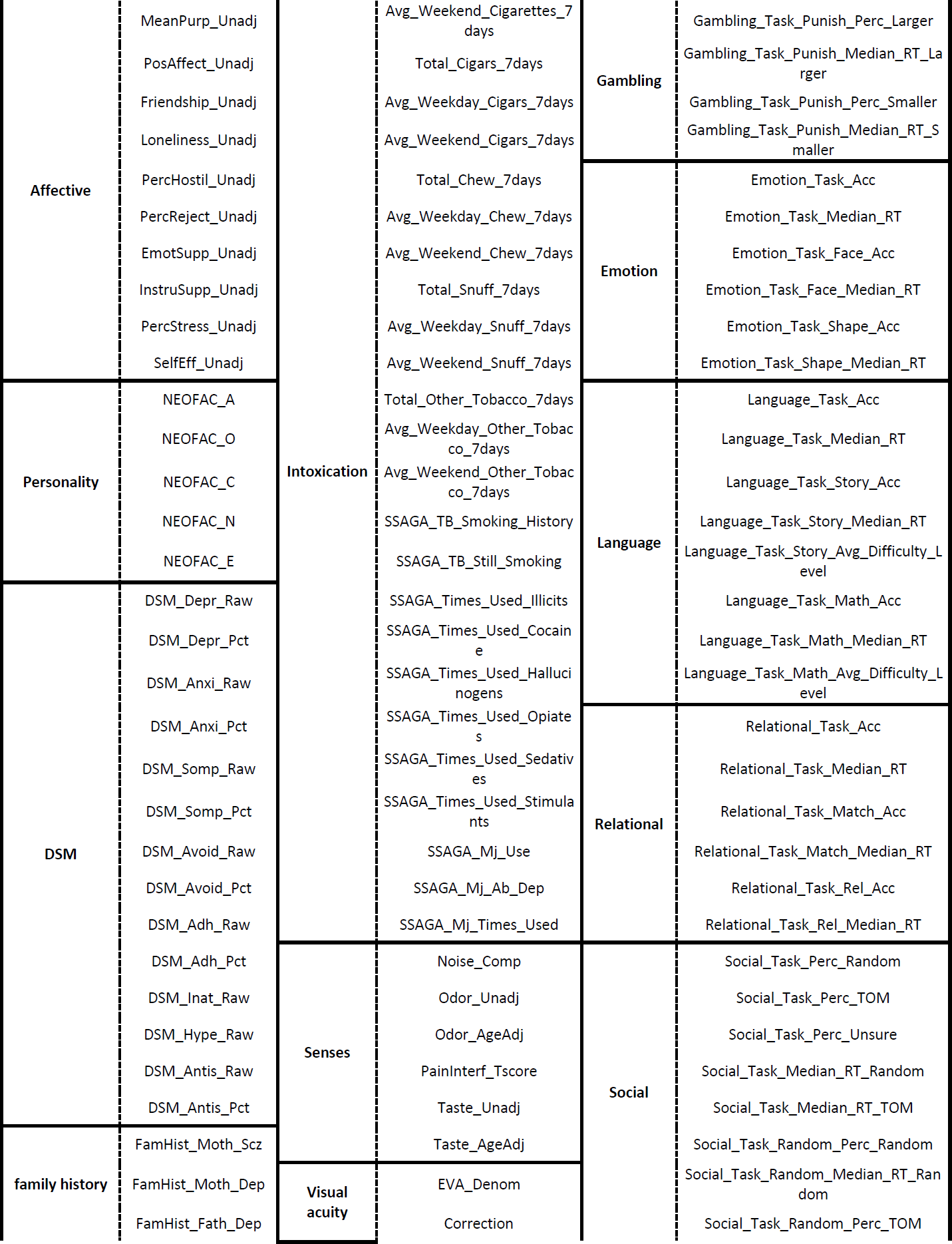
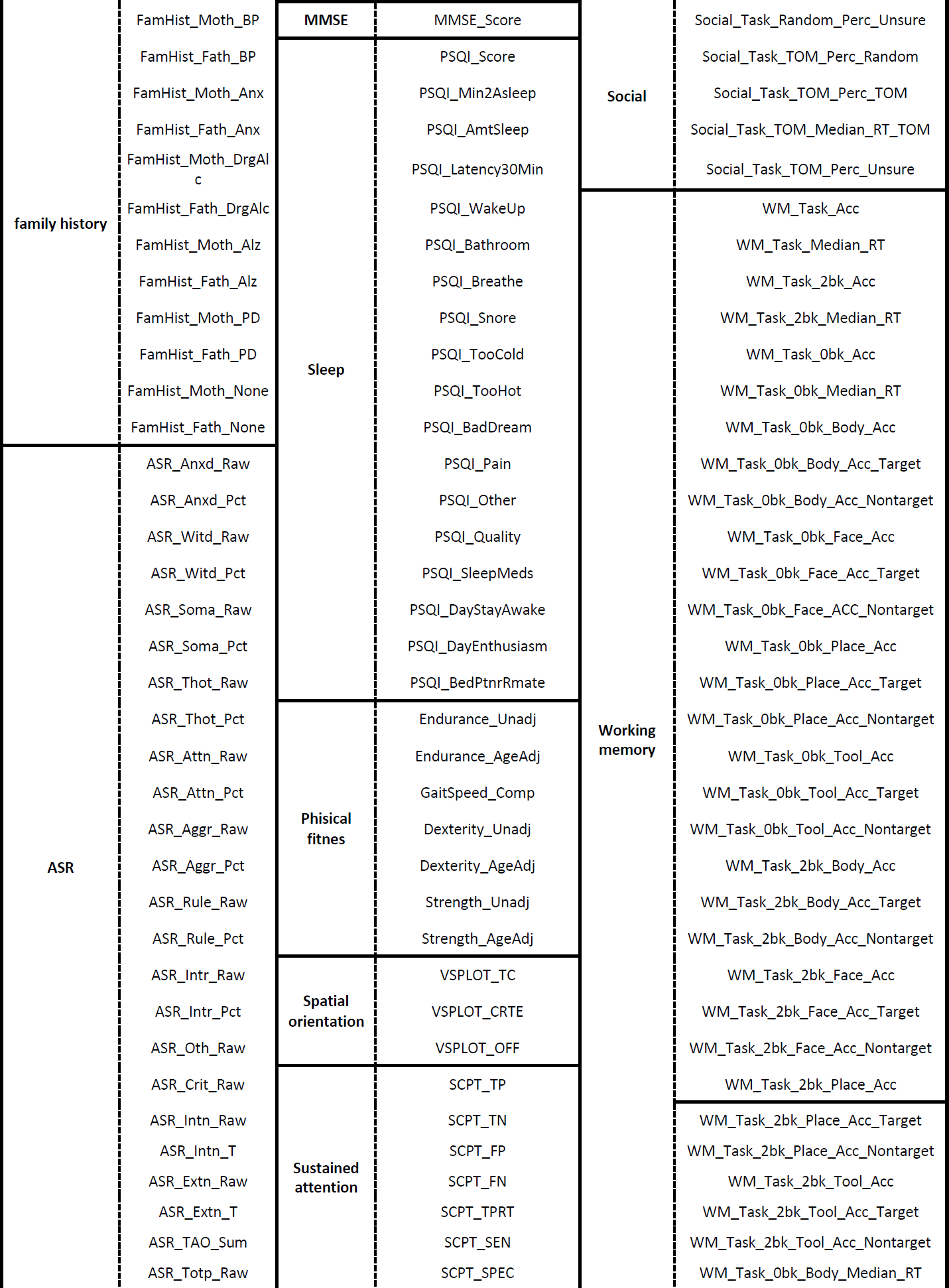
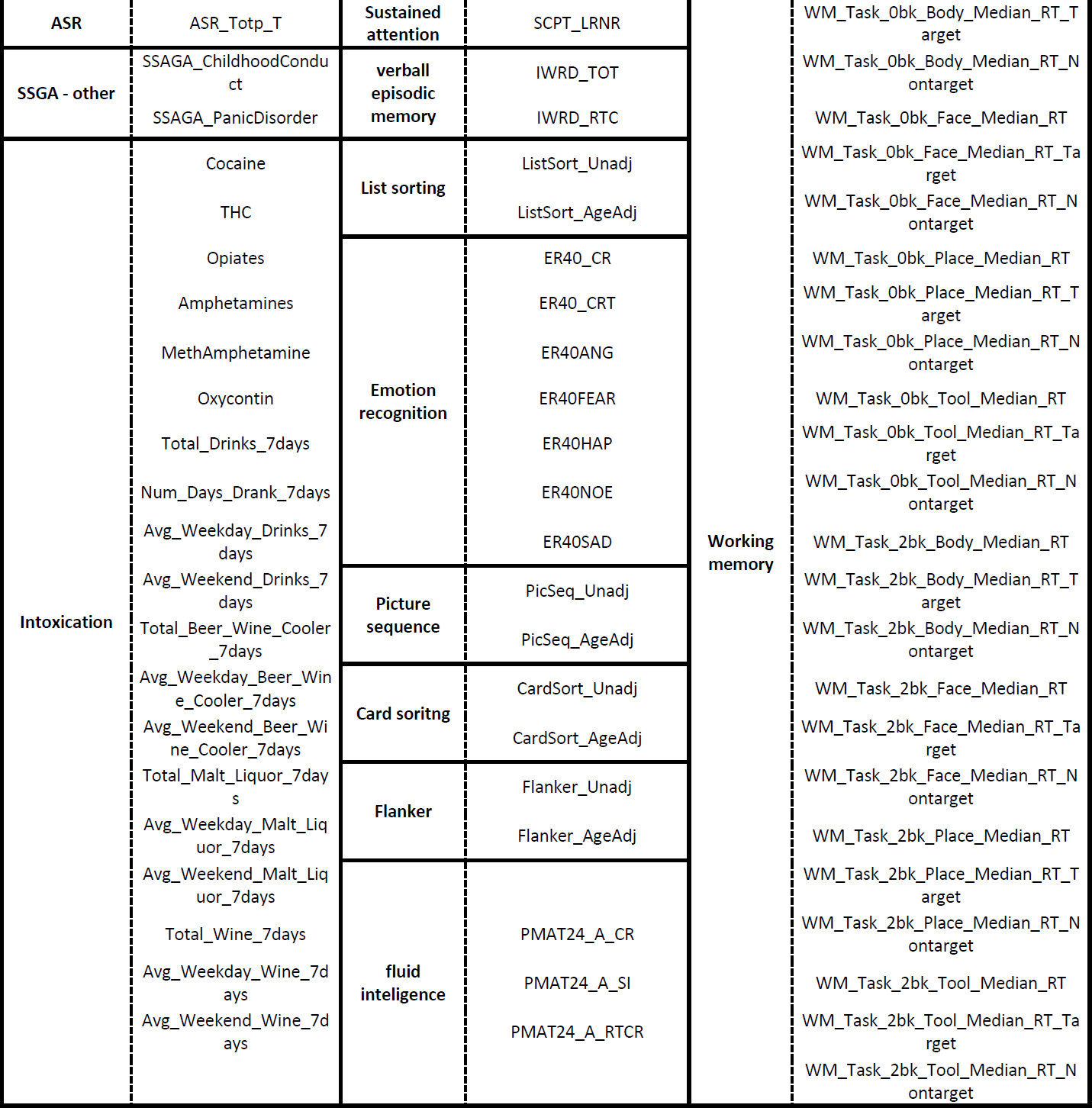
Summary of the behavioral/demographic measures present in the HCP sample.

### Individual features pre-processing

Prior to gray matter volume estimation, all participants’ T1 images were rigidly aligned using statistical parametric mapping software (SPM-12). Subsequently, images were segmented, normalized, and bias-field-corrected using ‘new segment’ from VBM-SPM12 (http://www.fil.ion.ucl.ac.uk/spm, ^22,23^), yielding images containing gray and white matter segments plus CSF. DARTEL ^35^ was then used to create a study-specific gray matter template to which all segmented images were normalized. Subsequently, all gray matter volumes were smoothed with a 9.4 mm FWHM Gaussian smoothing kernel (sigma=4 mm).

Structural MRI images were fed into FreeSurfer v5.3 software to extract measures for cortical thickness and areal expansion ^24,25^ (http://surfer.nmr.mgh.harvard.edu). The standard FreeSurfer preprocessing pipeline (recon-all) was applied to these images, in which a reconstruction of the cortical sheet was estimated using intensity and continuity information. Cortical thickness was determined as the closest distance from the gray/white boundary to the gray/cerebrospinal fluid (CSF) boundary at each vertex ^36^. Surface area in FreeSurfer is estimated as relative amount of expansion or compression at each vertex when registering each participant’s surface to a common atlas. Surface maps were resampled and mapped to a common coordinate system ^37^. During preprocessing, the data were registered onto the high-resolution average participant surface space (fsaverage), and a 10 mm FWHM surface-based smoothing kernel was applied.

Further, the Jacobian images for each subject are directly available from the HCP repository and the diffusion weighted data (DWI) was preprocessed using the DTIFIT routine from FSL ^27,38^ ((https://fsl.fmrib.ox.ac.uk/fsl) to create the FA, MO and MD images that were then feed into the TBSS pipeline ^26^.

Finally, for computational reasons ^13,14^, the VBM images were spatially down sampled to 4mm isotropic and the DTI images to 2mm isotropic voxels.

### Main results

In Table S2 we summarize the significant results obtained (FDR corrected q < 2.2×10^−4^). From left to right columns we present the component number, behavioral/demographical measure, correlation value, and the permutation p-value (PALM).

**Table S2:**
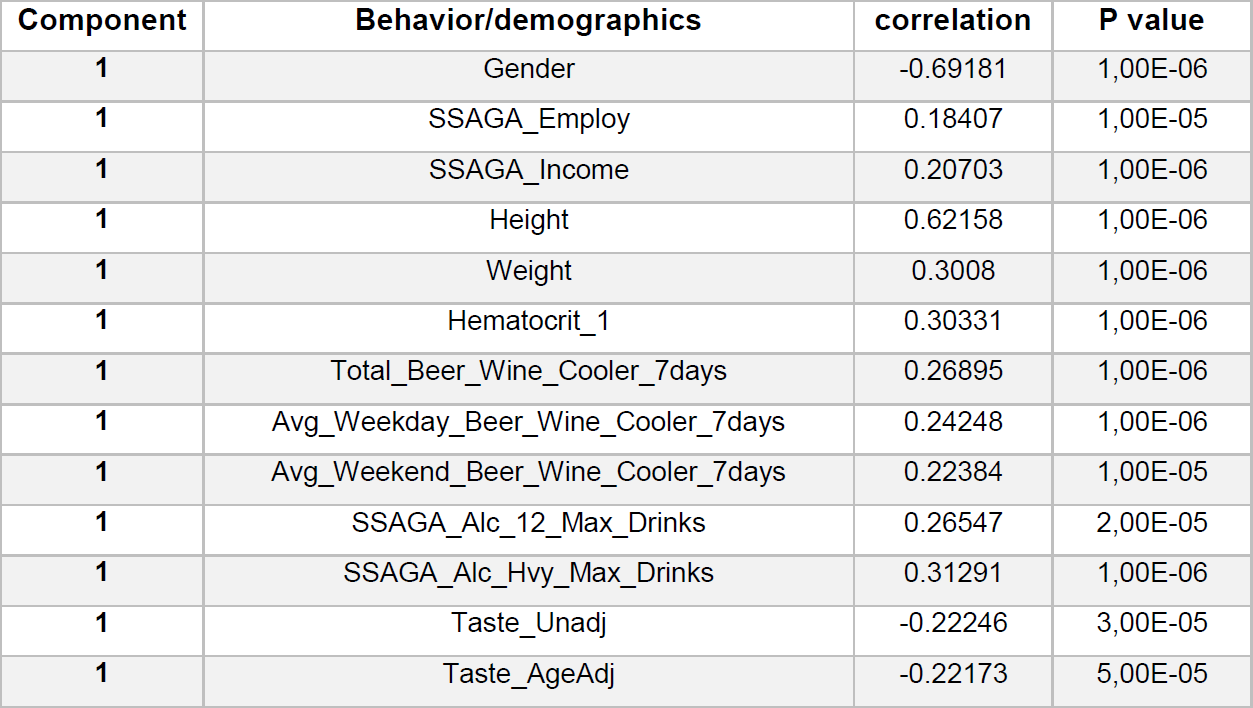
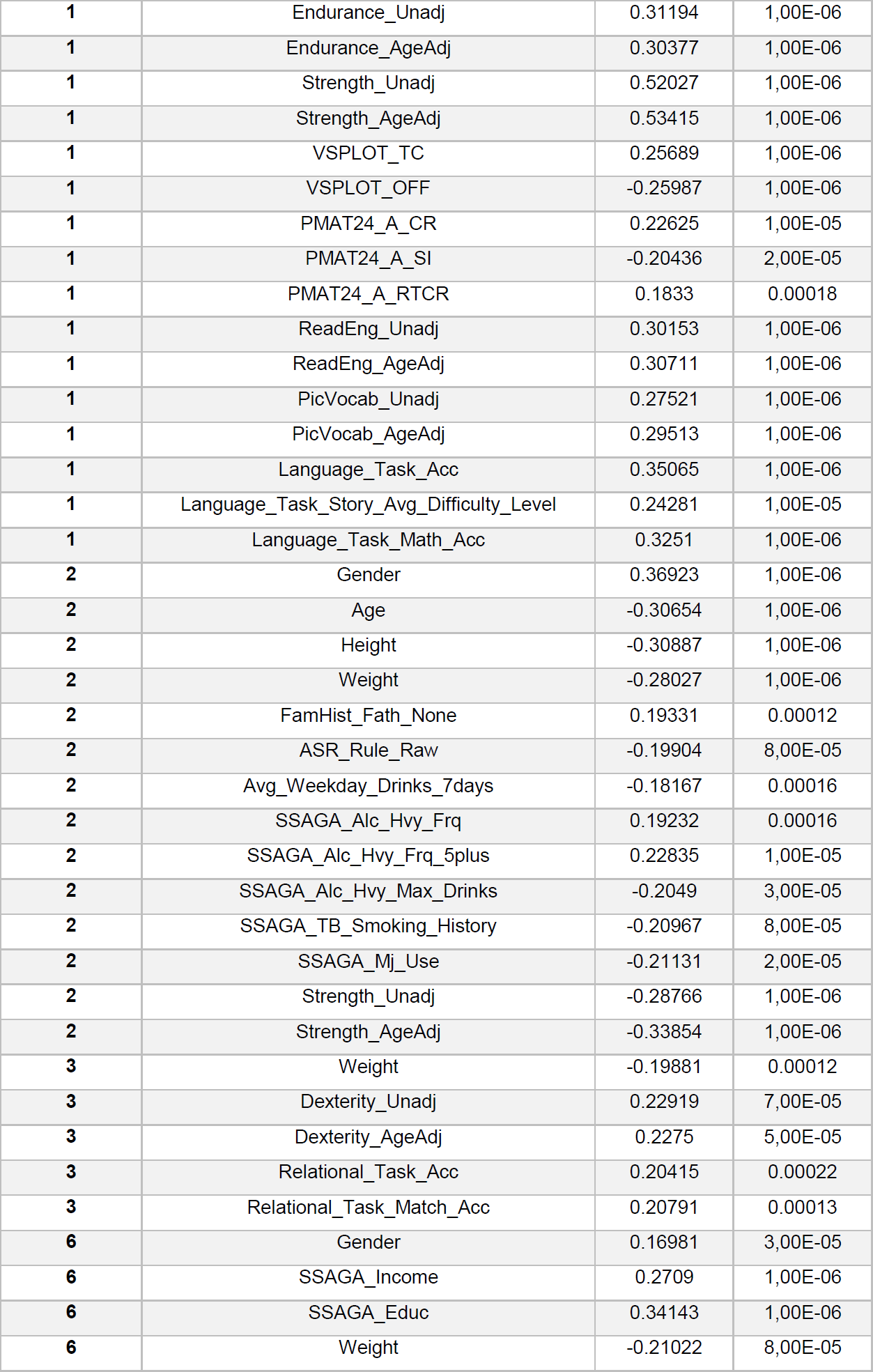
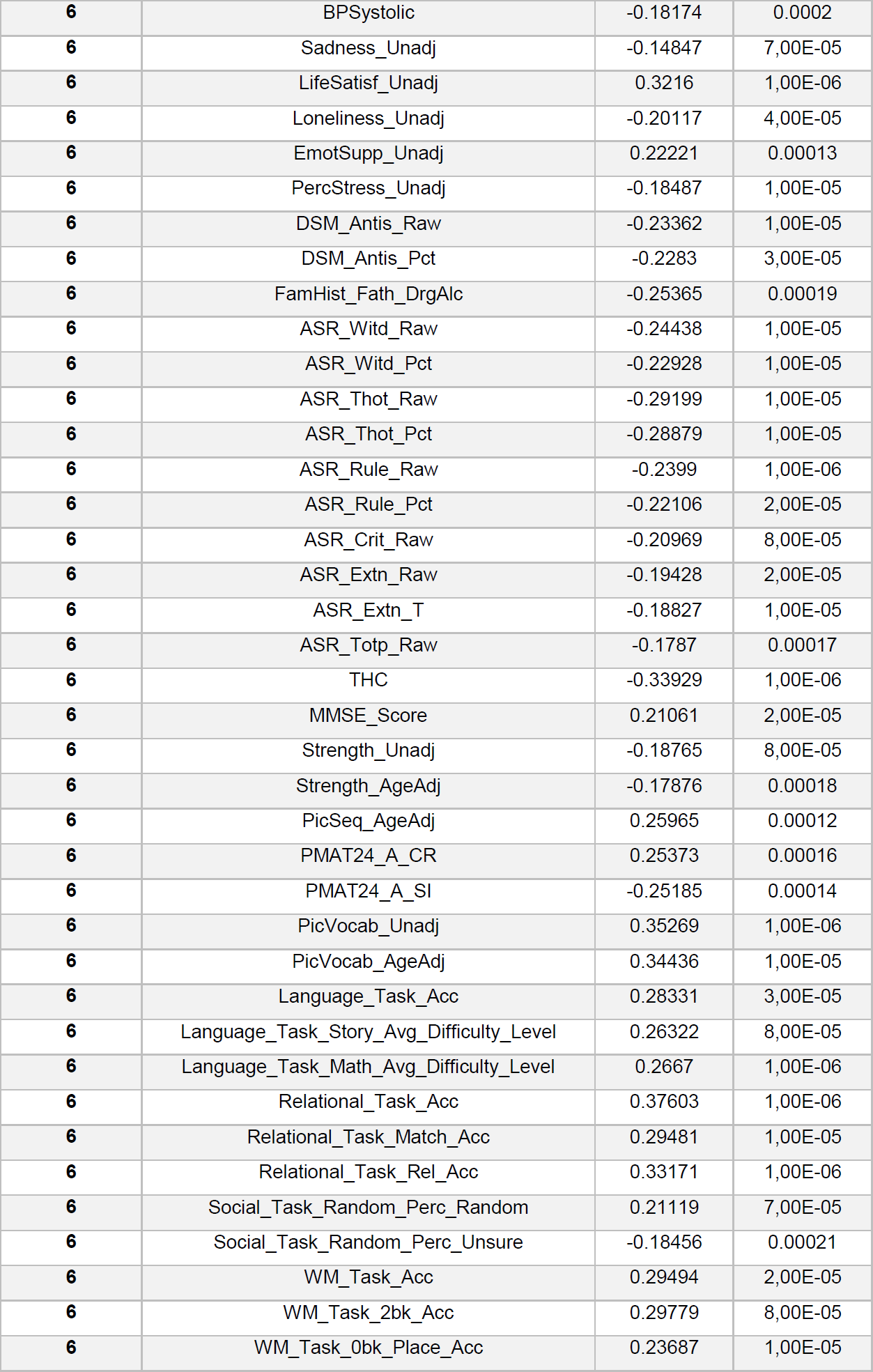
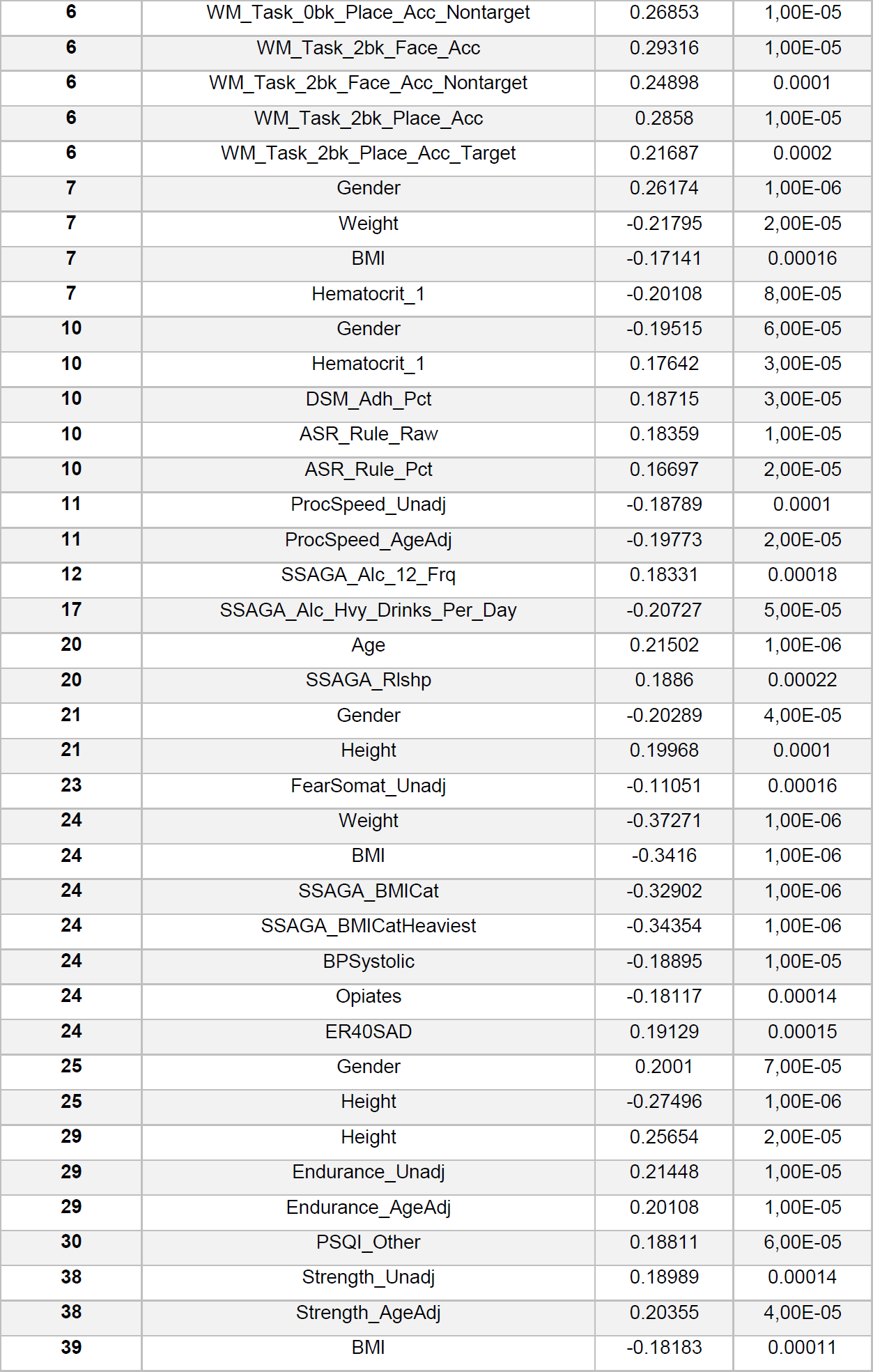
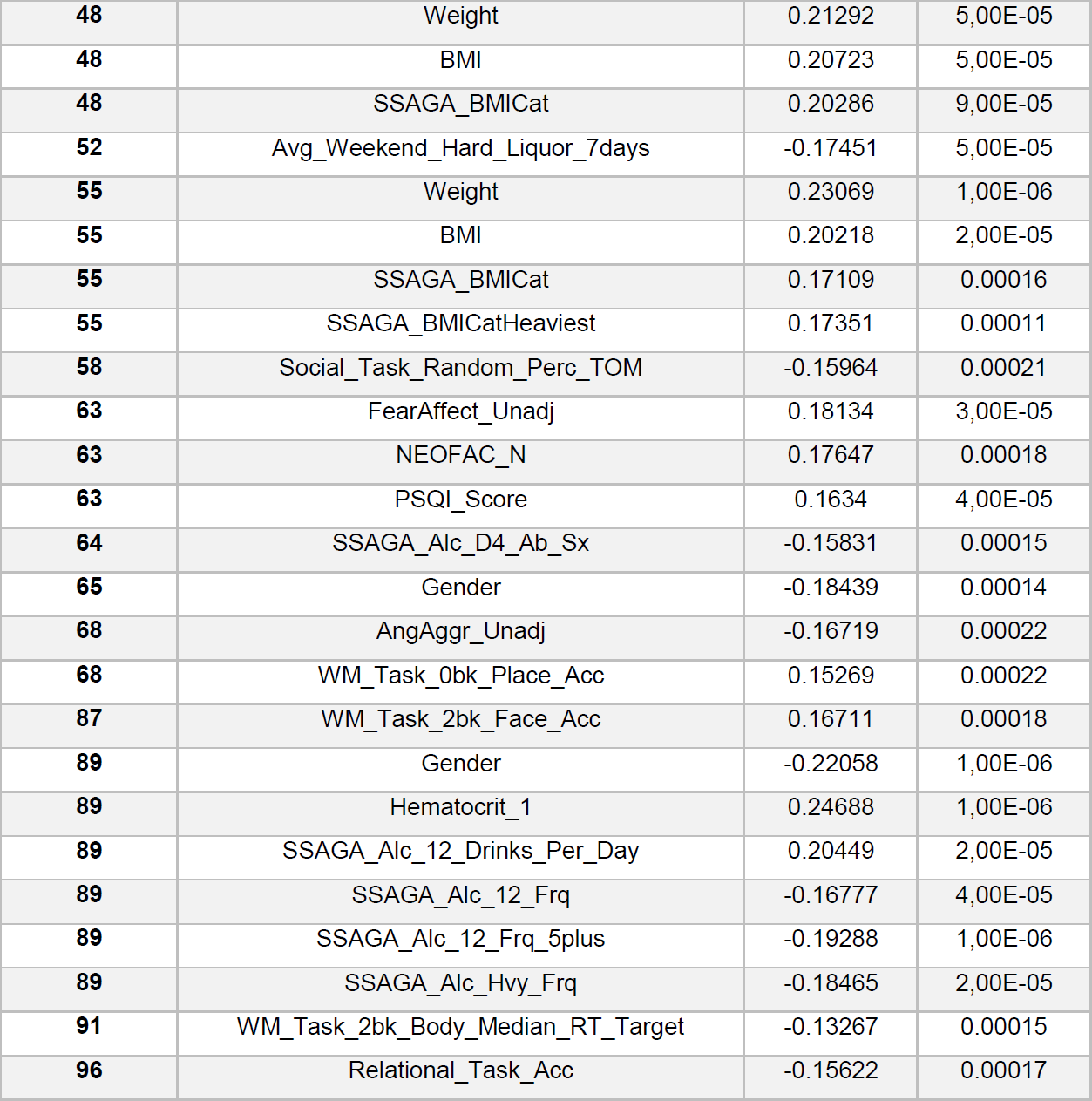
Significant results. First column presents the component number, second the behavioral or demographic measure it correlates with and third and fourth columns present the correlation value and the permutation p-value. Significance is defined at p<0.05 and we used FDR correction for multiple correction (q < 2.2×10^−4^).

### Spatial maps

In Figure S1 we report spatial maps associated with the components showing at least one Bonferroni corrected significant relationship with any behavioral measure (q < 1.4 x 10^−6^), which are not fully reported in the main text. The left column shows the independent component number. For components 1, 2 and 89 we show spatial maps for all modalities contributing to that component (Figure S2) i.e. VBM and PA for component number 1 and just VBM for components number 2 and 89. To provide a clearer interpretation we decided to show a selection of the relevant modalities for the other components and full nifty maps will be separately uploaded as SI. Here, we provide VBM, FA and CT for components number 7 and number 24; VBM and JD for component number 20 and finally VBM and FA for 25, 29 and 55.

**Figure S1:**
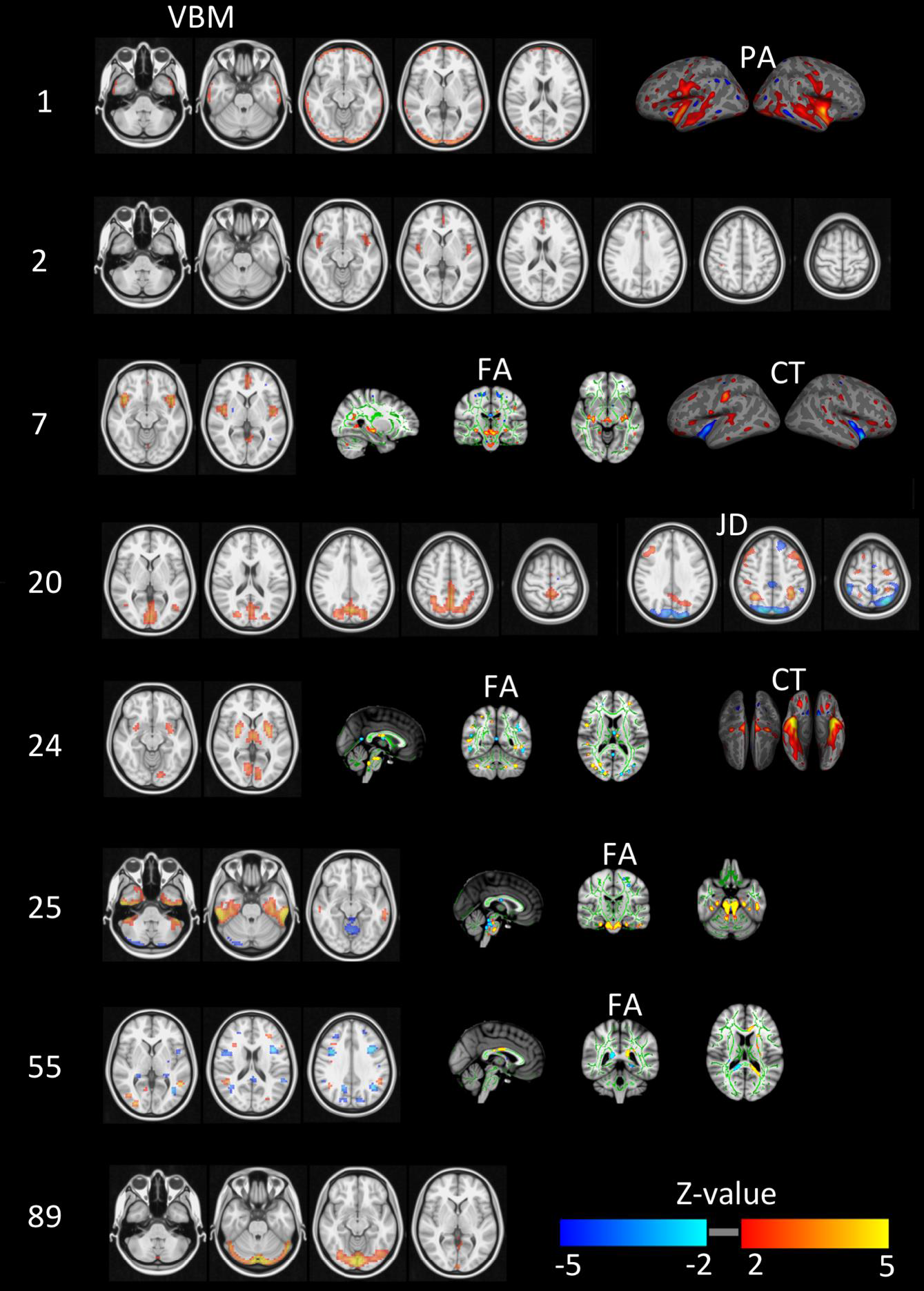
Summary of spatial maps associated with the components indicated in the most left column.

Component number 1 mainly relates to gender, physical strength and language and it is defined by voxel based morphometry and cortical area. Its associated spatial pattern appears to reflect brain size and cortical area differences in both temporal lobes. Component 2 is driven by VBM and maps into gender, age, height, weight and strength. Its spatial extent includes the paracingulate gyrus and bilateral insular and opercular cortex. Components 7 maps into gender and shows cingulate gyrus and insular cortex. Component 20 maps grey matter density in the posterior midline into age and relationship status. Components 24 maps into weight and body mass and its mapped into putamen, intracalcarine cortex and thalamus. Components 25 and 29 relate height with the inferior temporal gyrus and the cerebellum together with strong DWI weightings in the brainstem. Component 55 relates to weight and maps into the precentral gyrus and asymmetric differences in DWI measures. Component 89 maps into hematocrit and the lingual and occipital fusiform gyrus.

### Feature modalities relative contribution to components

In Figure S2 we color code the relative contribution of each feature modality to the components that show at least one Bonferroni corrected significant relationship to behavior or demographics measures (q < 1.4 x 10^−6^).

**Figure S2:**
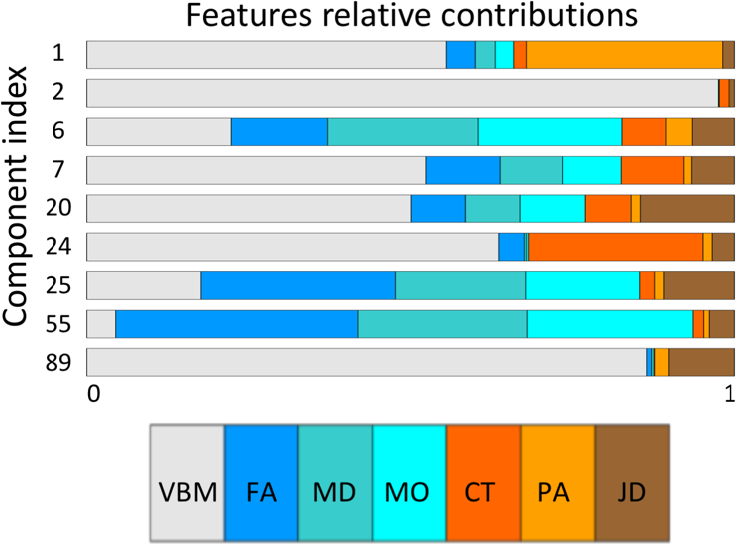
Relative contributions of each feature modality to the most relevant components.

### Robustness: model order

In this section we assess the robustness of the results with respect to the model order choice. To that end we perform a correlation analyses between the reported 100 dimensional factorization subjects-mode and that of a 90 and a 110 dimensional factorizations. In the top row of Figure S3 we present correlation matrices between a 100 dimensional factorization (y-axis) and a 90 and 110 dimensional factorizations (top left and right panels respectively). Only significant correlations after Bonferroni correction are reported (i.e. p-value smaller than 0.05 /(100×90) and 0.05/(100×110) respectively). For each component of the 100 dimensional factorization we present in the bottom row of Figure S3 the absolute correlation value with each of the components of the different dimensionality factorization. We appreciate that most components are recovered with high accuracy (r close to 1) independently of the order of the factorization. The black dashed lines represent the most relevant of the reported components, independent component number 6.

**Figure S3:**
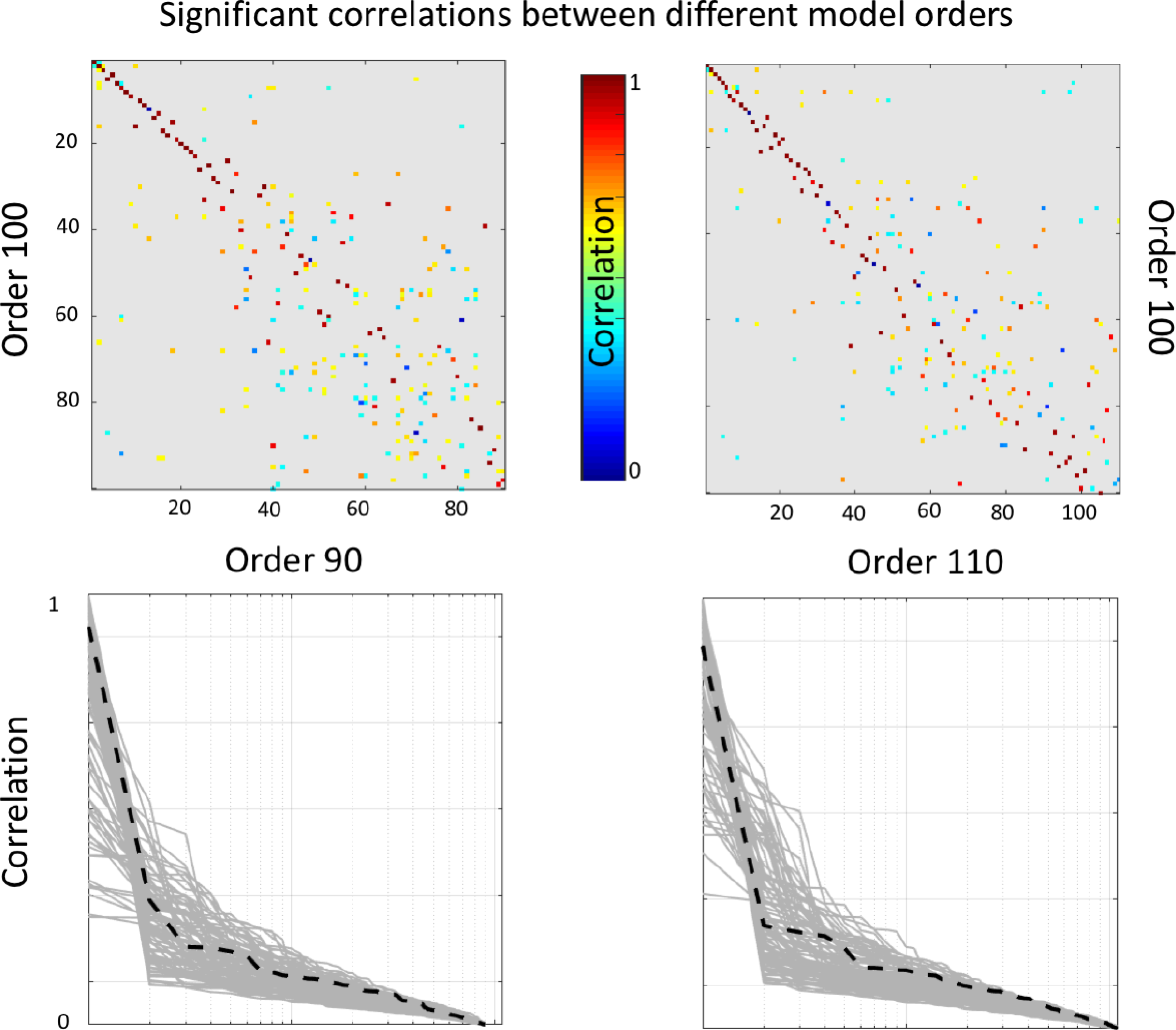
Top: Significant correlations between the reported (100 dimensional) factorization and a 90 dimensional (left panel) and 110 dimensional (right panel). Bottom: sorted absolute correlations for each of the components of the reported factorization with the other model orders components. The black discontinuous line represents component number 6.

### Robustness: analyses without the Jacobians

We present results summarizing the significant findings when performing an analogous analysis to the one reported in the main manuscript without the use of the JD feature. In Table S3 we present a comparison between the positive-negative mode as reported in the main text, and the set of behavioral measures significantly associated to component number 9 on the new analysis.

**Table S3:** Comparison between the positive-negative mode presented in the main text and the multi-modal analyses excluding the JD feature (right column).

| Behavioral and demographics | Linked ICA component 6 - correlations | Linked ICA (no JD) - correlations |
| --- | --- | --- |
| Relational Task | 0,35 | 0,29 |
| Picture Vocabulary test | 0,35 | 0,29 |
| Life Satisfaction | 0,32 | 0,3 |
| Years of Education and Income | 0,3 | 0,3 |
| Working Memory | 0,27 | 0,27 |
| Language | 0,27 | 0,26 |
| Picture Sequence | 0,26 | 0,23 |
| Fluid intelligence (correct responses) | 0,25 | 0,24 |
| Emotional Support | 0,22 |  |
| Social Task (Random Perc. Random) | 0,21 |  |
| MMSE_Score | 0,21 |  |
| Gender | 0,16 |  |
| Sadness | -0,15 |  |
| Perceived Stress | -0,18 |  |
| Social Task (Random Perc. Unsure) | -0,18 |  |
| BPSystolic | -0,18 |  |
| Weight and Strength | -0,19 | -0,23 |
| Loneliness | -0,20 |  |
| DSM | -0,23 | -0,25 |
| ASR | -0,23 | -0,24 |
| Family History Drugs & Alcohol | -0,25 |  |
| Fluid intelligence (skipped responses) | -0,25 | -0,25 |
| Positive test for THC (cannabis) | -0,34 | -0,31 |

Note that component number 9 in this analysis (without the JD feature) corresponds to the component number 6 reported in the main text and it recovers the strongest behavioral associations. Further, this component presented other significant relationships not appearing significant in the main reported mode, e.g. personality related measures (NEOFAC-C).

### Robustness: Analyzing morphometric differences

In this section we perform an independent component analyses (ICA) factorization^29^ only of the Jacobian determinant matrices followed by post-hoc correlation analyses with the behavioral and demographic measures. We performed a 100-dimensional factorization and found a set of 9 components significantly correlating (Bonferroni corrected) with the component number 6 obtained from the multi-modal Linked ICA analyses. In Table S4 we present the correspondence between the positive-negative mode we found through the multimodal Linked ICA analyses and these 9 components. As before, for the multimodal analyses we only report correlations to behavior significant after FDR correction and for the JD analyses we mark with double asterisk the significant relationships after FDR correction and with a single asterisk the nominal or uncorrected significant relations (p<0.01). We appreciate that although sub FDR significance threshold we observe some correspondence between the purely morphometric differences and the positive-negative mode, these relationships disappear after statistical correction for multiple comparisons.

**Table S4:** Summary of the uni-modal analyses using the JD feature. In the second row all relationships are significant after multiple comparison correction. For the uni-modal analysis (the third row), significant associations after multiple comparison correction are denoted with a double asterisk and nominal significant but not significant after multiple comparison correction are marked with a single asterisk.

| Behavioral and demographics | Linked ICA component 6 - correlations | ICA in JD (9 components) - correlations |
| --- | --- | --- |
| Relational Task | 0,35 | 0,17* |
| Picture Vocabulary test | 0,35 | 0,23 |
| Life Satisfaction | 0,32 | 0,15* |
| Years of Education and Income | 0,3 | 0,17 |
| Working Memory | 0,27 | 0,2 |
| Language | 0,27 | 0,17* |
| Picture Sequence | 0,26 |  |
| Fluid intelligence (correct responses) | 0,25 | 0,2* |
| Emotional Support | 0,22 | 0,13* |
| Social Task (Random Perc, Random) | 0,21 | 0,14* |
| MMSE_Score | 0,21 | 0,12* |
| Gender | 0,16 | 0,14* |
| Sadness | -0,15 | -0,14* |
| Perceived Stress | -0,18 | -0,15* |
| **Social Task (Random Perc. Unsure)** | -0,18 | -0,11* |
| **BPSystolic** | -0,18 | -0,12* |
| **Weight and Strength** | -0,19 | -0,15* |
| **Loneliness** | -0,20 |  |
| **DSM** | -0,23 | -0,09* |
| **ASR** | -0,23 | -0,17* |
| **Family History Drugs & Alcohol** | -0,25 | -0,17* |
| **Fluid intelligence (skipped responses)** | -0,25 | -0,22* |
| **Positive test for THC (cannabis)** | -0,34 | -0,17* |

